# A validated and interpretable predictive model of cruzain inhibitors

**DOI:** 10.1101/2020.06.02.129411

**Authors:** Jose G. Rosas-Jimenez, Marco A. Garcia-Revilla, Abraham Madariaga-Mazon, Karina Martinez-Mayorga

## Abstract

Chagas disease affects 8–11 million people worldwide, most of them living in Latin America. Moreover, migratory phenomenon have spread the infection beyond endemic areas. Efforts for the development of new pharmacological therapies are paramount, as the pharmacological profile of the two marketed drugs currently available, nifurtimox and benznidazole, needs to be improved. Cruzain, a parasitic cysteine protease, is one of the most attractive biological targets due to its roles in parasite survival and immune evasion. In this work, we generated Quantitative Structure-Activity Relationship linear models for the prediction of pIC_50_ values of cruzain inhibitors. The statistical parameters for internal and external validation indicate high predictability with a cross-validated correlation coefficient of 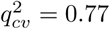 and an external correlation coefficient of 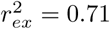. The applicability domain is quantitatively defined, according to QSAR good practices, using the leverage method. A qualitative interpretation of the model is provided based on protein-ligand interactions obtained from docking studies and structural information codified in the molecular descriptors relevant to the QSAR model. The model described in this work will be valuable for the discovery of novel cruzain inhibitors.

**Author summary:** Chagas disease is a major health problem in Latin America. The disease involves a long-lasting silent phase that usually culminates in serious or fatal heart damage. Despite its prevalence, there are only two antichagas approved drugs available. Despite these drugs have been in the market for more than 50 years, significant undesirable side effects and modest effectiveness in the chronic phase are prevalent. The need of new drugs to treat this disease is evident. Cruzain is a vital protein for the survival of *Trypanosoma cruzi,* the parasite causative of Chagas disease. Inhibition of this species-specific protein has been associated with improvements in pharmacological effects in animal models. Thus, blocking the activity of cruzain is an attractive approach for the development of antichagas agents. In this work, we present a validated mathematical model capable of predicting the cruzain inhibition value of a molecule from its chemical structure. This model can contribute to the identification of potential pharmacological alternatives against Chagas disease.

## Introduction

Chagas Disease affects 8-11 million people in 21 Latin American countries, there is an estimation of 70 – 150 million people at risk of infection [1,2]. Migration phenomenon have contributed to the spread of the parasite into non-endemic areas such as the United States, Europe, New Zealand, and Australia [1]. Chagas disease is a vector-borne parasitic infection caused by *Trypanosoma cruzi* and it is transmitted by the three main genera of triatomine bug, *Triatoma, Rhodnius,* and *Panstrongylus.* World Health Organization has recognized this infection as a Neglected Tropical Disease (NTD) because of its persistence in developing countries, being a major economic and social problem in these regions, and one of the main causes of premature death for heart failure [2–4]. It was previously reported that this disease causes an estimated loss of 752 000 working days in southern American countries [4], which implies an economic burden of about US$1.2 billion in productivity. Globally, this parasitic infection has an estimated annual cost of $627.46 million, and 10% of this affects non-endemic countries [4]. Currently, there are only two approved drugs for the treatment of Chagas Disease: Nifurtimox (NFX) and Benznidazole (BZ). Both NFX and BZ have similar efficacy during the acute phase of infection, with 88 – 100 % of negative parasite detection after treatment with NFX and up to 80 % for BZ [5]. However, in the chronic phase, the rate of negative tests for the disease after treatment falls to 7 – 8 % [5], and there are significant side effects, including anorexia, weight loss, paresthesia, nausea, and vomiting, among others [3,5]. Recent therapeutic research is focused on specific biological targets, which include cysteine proteases, enzymes in trypanothione metabolism, enzymes in ergosterol biosynthesis and the kinetoplastid proteasome [5].

Cruzain is a cathepsin L-like cysteine protease present in all stages of the parasite life cycle. It plays significant roles in the trypanosomal growth, survival and evasion from the host immune response. Plasma membrane-anchored cruzain degrades the Fc fraction of antibodies, overcoming the classic path of complement activation [3,6]. In the amastigotic intracellular stage, this cysteine protease degrades transcription factors, such as NFkB and thus prevents the activation of macrophages [3]. Cruzain generates the bloodstream pro-inflammatory peptide Lys-bradykinin, which activates host immune cells, promoting the parasite uptake and spread by phagocytosis [6]. The use of cruzain inhibitors in animal models has shown to be effective in clearing the parasite burden, even in the chronic phase. The vinyl-sulphonic compound known as K777 was one the first proof-of-concept studies about anti-tripanosomal activity of cruzain inhibitors in animal models [7,8]. Parasite death induced by cruzain inhibitors is attributed to the accumulation of a peptide precursor in the Golgi complex. Therefore, these *in vitro* and *in vivo* evidence have validated cruzain as a potential biological target for Chagas Disease [3,6]. A variety of chemotypes for cruzain inhibition have been explored through Structure-Activity Relationships (SAR) analysis, high-throughput screening and docking methods. The most potent molecules belong to the vinyl-sulfone derivatives, oxadiazoles, nitrile-containing peptidomimetics, and thiosemicarbazones, with a broad range of biological activities among chemical families [2]. These molecules should be further optimized by increasing their selectivity towards parasite vs human cathepsins, and they should be neutral at physiological pH, to avoid concentration in lysosomes and off-target effects [2].

Quantitative Structure-Activity Relationships (QSAR) is a ligand-based approach that mathematically correlates structural properties of molecules with their biological activity. QSAR modeling is widely used in drug discovery, especially in the prediction of enzyme inhibition and ADME-Tox properties [9]. In virtual screening, validated QSAR models are used for prioritizing molecules for experimental evaluation. Carefully validated QSAR models have rendered novel chemotypes and scaffolds with a desirable biological activity [10]. The quality of a QSAR model can be evaluated using the OECD principles [11]. These principles are a series of guidelines originally developed for the use of QSAR modelling for regulatory purposes, but they became a valuable tool in the standard QSAR practice [11,12]. In this work, we explored public databases of structurally diverse cruzain inhibitors for the generation of QSAR predictive models of this biological endpoint. The structural properties, encoded by molecular descriptors, are rationalized in terms of protein-inhibitor interactions, using molecular docking, thus providing a possible mechanistic interpretation of the model. This work will be useful in the search of cruzain inhibitors.

## Materials and methods

### Data compilation, curation, and pre-processing

Cruzain inhibitors were collected from the ChEMBL (release 24) database, searching by molecular target using the keyword *cruzain.* Molecules annotated with IC50 values were selected and duplicated or missing values compounds were eliminated. Finally, a selection based on the same experimental protocol for IC_50_ determination was performed. The selected experimental procedure is a competitive fluorescence assay in the presence of detergent, as reported by Babaoglu *et al* [13]. The detergent is used to avoid aggregation, which is the main cause of false positives in exploratory and high throughput screens [2, 14]. Structural and biological information of the compounds was verified in the corresponding original publications, and when required, the discrepancies were fixed. IC_50_ values were transformed to pIC_50_. The final dataset consisted of 110 structurally diverse cruzain inhibitors. The 2D and 3D coordinates of these molecules were calculated from their SMILES representation, using the *wash* tool in the software Molecular Operating Environment 2019.01 (MOE) [15]. Lastly, the structures were energy minimized using the MMFF94x force field. The curated database is available in S1 Table.

To summarize the chemical diversity in the set, the MACCSKeys fingerprints as implemented in RDKit [16] package were calculated for every molecule. A clustering calculation was performed using the affinity propagation algorithm, with the Tanimoto similarity matrix as affinity measure. The chemical structures for the representative molecules in every cluster are presented below.

### QSAR modeling

#### Descriptor calculation and feature selection

All molecular descriptors available in MOE were calculated, including topostructural and topochemical indices, subdivided Van der Waals surface areas, VolSurf potentials, and physicochemical properties such as dipolar and hydrophobic moments. The dataset was randomly split into a training set (88 molecules, 80%) and a test set (22 molecules, 20%). Descriptors were scaled to [0,1] range using Eq (1) and those in the test set were scaled according to the training set. Constant descriptors (zero variance) were filtered out.

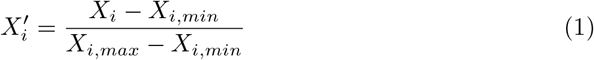

Feature selection and model calculation were performed in Weka 3.8 [17,18]. Selection of relevant features was carried out using the Correlation-Based Feature Selection (CFS) with a Greedy Stepwise algorithm [19]. Briefly, CFS calculates a merit score, *M_s_*, on a subset of variables through Eq (2), where *r_fc_* is the average pair-wise correlation coefficient between features and the dependent variable, *r_ff_* is the average pair-wise correlation coefficient between features themselves, and *k* is the number of features. Higher merit values involve a higher correlation with the dependent variable and a less correlation between features, penalizing high-dimensional sets also. In the Greedy Stepwise algorithm, the variables are sequentially added until the merit reaches a maximum.

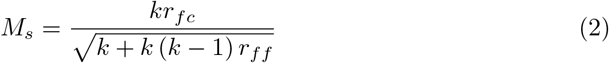

The subset of features with the highest score were used in the generation of the Multiple Linear Regression model, as implemented in Weka 3.8.

#### Model validation

The goodness of fit for the model was estimated by calculating the following statistical parameters: coefficient of determination (*R*^2^), adjusted coefficient of determination (*R*^2^ – *adj*), *F* statistic (variance ratio) and its associated p-value. Internal validation was carried out through the *k*-fold leave-some-out cross-validation with *k* = 10. The cross-validated correlation coefficient 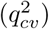 is reported to evaluate the robustness of the model. Model predictability was assessed by applying the generated equation to calculate the biological data of the test set. Using these results, the Golbraikh and Tropsha (G&T) external validation parameters were calculated in the Enalos nodes for Knime [20–22]. Golbraikh and Tropsha parameters use regression through the origin to estimate the deviation of the model with respect to the ideal QSAR regression. Basically, these parameters compare the differences of the coefficients of determination and slopes of the fitted model and the regression forced to the origin, *R*_0_ and *k*. The model is considered predictive if all parameters are within defined thresholds [20].

#### Applicability Domain

The predictivity of a QSAR model is framed by the nature of the molecules in the training set. The applicability domain is the quantitative delimitation of the chemical space where predictions are reliable. In this work, the applicability domain was defined using the leverage method [23]. Leverage values, *h_i_*, are computed using Eq (3), where **X** is the descriptor matrix of the training set and *x_i_* is the descriptor vector for a query molecule.

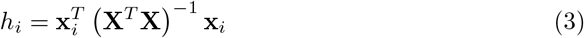

Basically, leverage values are proportional to the distance of the molecule from the centroid of the training set. Thus, compounds above a threshold are far from the explored chemical space and therefore, their predicted biological activity will be unreliable. Typically, the threshold, *h_max_*, is computed with Eq (4), where *p* is the number of features and *n* is the number of molecules in the training set.

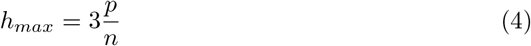

Leverage and limit values were computed with the Applicability Domain node calculator of Enalos for Knime [24,25]. Results are presented in a Williams plot (leverage vs standarized residuals), where outliers in the activity domain or structurally influential, can be visually detected. The Williams plot is a representation of the chemical space spanned by the model.

### Docking calculation

The coordinates of cruzain were downloaded from the Protein Data Bank with PDB-ID code 3KKU [26]. This structure is reported with a resolution of 1.28 Å, and it is co-crystalized with a benzimidazole derived, a non-covalent ligand. The protein was prepared in MOE as follows: hydrogen atoms were added according to protonation states at pH 7.0 and Gasteiger-Marsili charges were computed [27]. The protein structure file was converted to the PDBQT format. Gasteiger charges were also computed for the ligands and they were converted to PDBQT format. The docking calculation was performed in AutoDock Vina [28]. The search space was extended in the binding site of the cruzain with a box of size 24 Å * 30 Å * 20 Å. The docking calculation was performed in 10 repetitions, and the conformations with the best score per molecule were selected to generate a database of bound conformations. These data were used to generate protein-ligand interaction fingerprints in MOE.

The similarity maps tool included in the RDKit module for python was used to generate partial charges and SLogP diagrams. These diagrams show the atomic contributions to logP, calculated with the Wildman and Crippen algorithm [29], and the Gasteiger partial charges [27]. The 2D depictions of molecules in the similarity maps were generated by projecting the calculated 3D conformation of the molecules, so that these depictions resemble their docked pose.

## Results and Discussion

In this work, we present the preparation and analysis of a data set of 110 cruzain inhibitors annotated with pIC_50_ values. The distributions of the biological activity values of the training and test sets are shown in Fig 1. The pIC_50_ values ranges from 3.48 to 10.0 units, from nanomolar to sub-millimolar scale. Typically, the reported experimental error for this biological assay is around 2.0 *μM*, which implies that the range of pIC_50_ values of this dataset is more than five times higher than the experimental error, in agreement with general recommendations and best practices for QSAR modeling [10]. Noteworthy, the biological activity values of the test set lie within that of the training set as shown in the histogram of the Fig 1 and with no gaps within bins.

**Fig 1.**
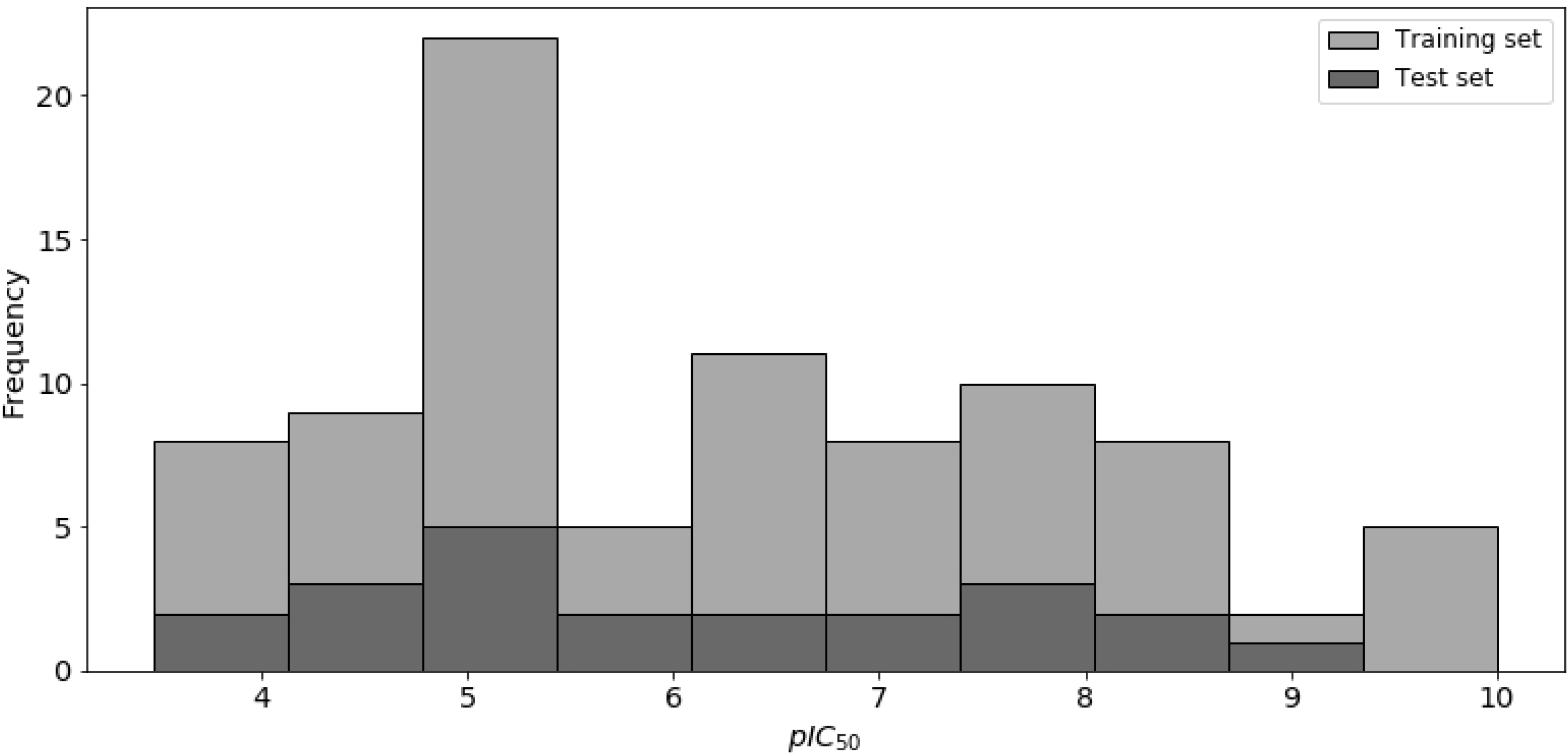
Distribution of pIC_50_ values of 110 cruzain inhibitors. Molecules in the training set (88) are shown in gray, and 22 molecules in the test set are shown in black. The inhibitory potency of the test set fall within interval of pIC_50_ values of the training set.

To summarize the chemical diversity contained in the dataset, we performed a clustering analysis with the affinity propagation algorithm, using Tanimoto similarity from MACCSKeys fingerprints as affinity measure, in the RDKit and Sci-Kit learn modules in Python. Fig 2 depicts the calculated clusters, along with the representative structure from every cluster. The chemical families include thiosemicarbazones, acylhydrazines, oxadiazoles and nitrile-containing peptidomimetics. Thiosemicarbazones are the most numerous compounds in the set and peptidomimetics are the most potent known inhibitors, as has been reported previously [2].

**Fig 2.**
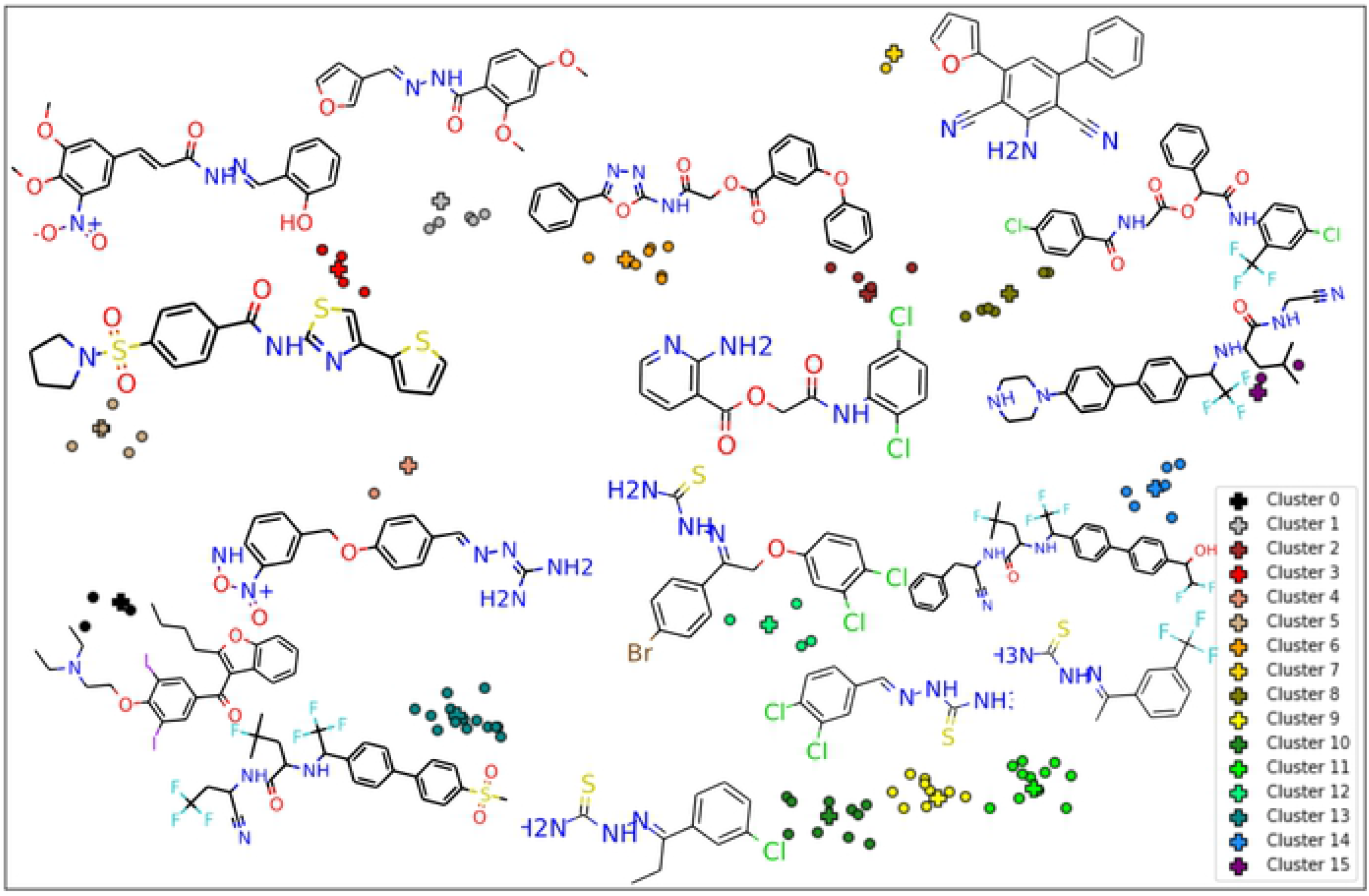
Representation of the molecular diversity in the dataset. Molecules are grouped according to the clustering results by affinity propagation. Structures for representative molecules, highlighted with a cross-shaped mark, are shown for every cluster.

The MLR algorithm generates an explicit equation, consisting of a linear combination of molecular descriptors. After selecting the feature subset which renders the maximum merit score, as described in the methods section, Eq 5 was obtained for the estimation of pIC_50_ values. A brief definition of the descriptors involved in the model is presented in Table 1. In general, these descriptors account for electrostatic (E_ele_ and PEOE descriptors), hydrophobic (SLOGP and vsurf_ID8 descriptors) and hydrophilic (vsurf_W related descriptors) properties. These features are crucial for the establishment of potential intermolecular interactions required for the binding of ligands into the active site. Therefore, the linear equation may be related to the presence of such features in the binding process.

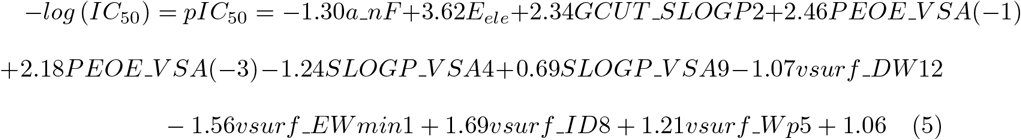

**Table 1.**
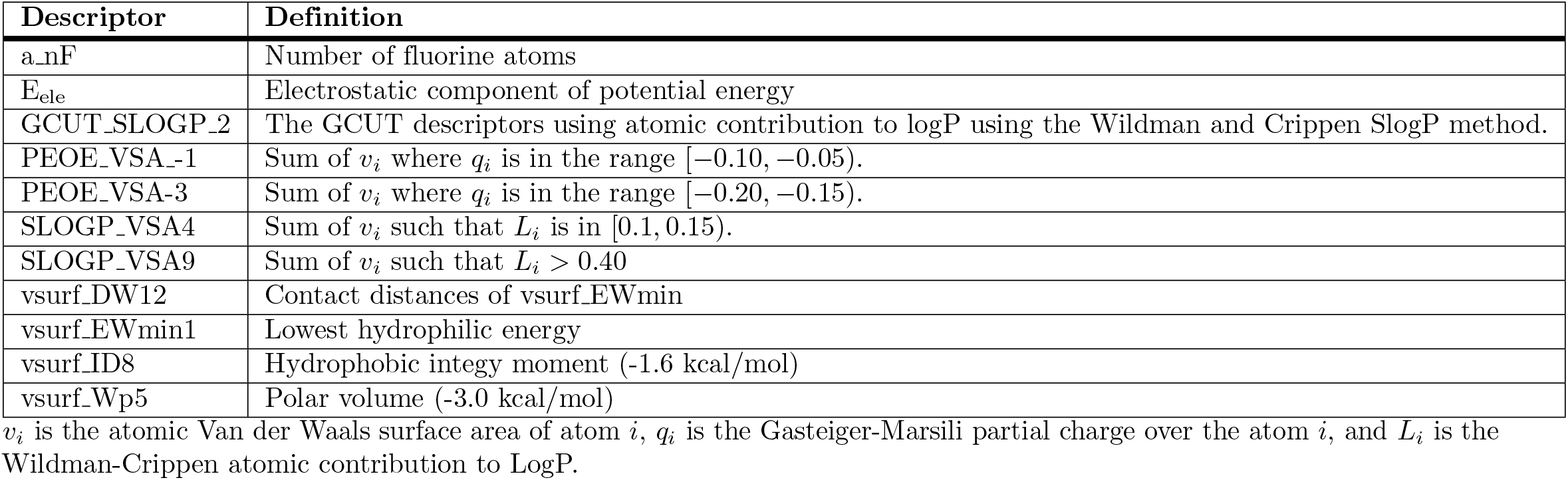
Definition of molecular descriptors selected in the linear equation of the model.

Statistical parameters describing the goodness of fit for the model are presented in Table 2. Coefficients of determination near to 1 indicates that a high ratio of variance present in the original data is explained by the model. In this case, 83% of the variance already present in the pIC_50_ of the training set is explained by Eq 5. The ratio of the mean squared error of the one-parameter model and the generated model is measured by the *F* statistic. If this ratio is high enough, the prediction made by Eq 5 has an error less than the native variability in the data. The *F* value for this model is presented in Table 2 along with its associated *p* — *value*. Making the assumptions of the linear model, the probability of finding an *F* ratio of 34.08 or higher, for a 10 parameter equation, is less than 0.001, if the model error is equal to the variability in data. Given this low probability value, this hypothesis can be rejected and accept that predictions made by the model equation are more accurate than just the mean value around the standard deviation for the original data.

**Table 2.** Statistical parameters describing the goodness of fit for the model.

| Parameter | Value |
| --- | --- |
| $R^2$ | 0.83 |
| $R^2$-adjusted | 0.81 |
| $F$ ratio | 34.08 |
| $p$-value | < 0.001 |

Since conclusions derived from statistical parameters rely on the parametric assumptions, their fulfillment were tested by means of an analysis of residuals, shown in Fig (3). In the linear model, the dependent variable, *Y_i_*, has a normal distribution around the predicted value *Y_i_*, thus the prediction error, *Ŷ_i_* — *Ŷ_i_*, must follow a normal distribution with a mean of 0. The lower panel of Fig (3) shows the quantile plot for the calculated errors and the theoretical normal distribution. Most of the values in the quantile plot follow a straight line, suggesting a very near behavior to a normal distribution, achieving the normality requirement.

**Fig 3.**
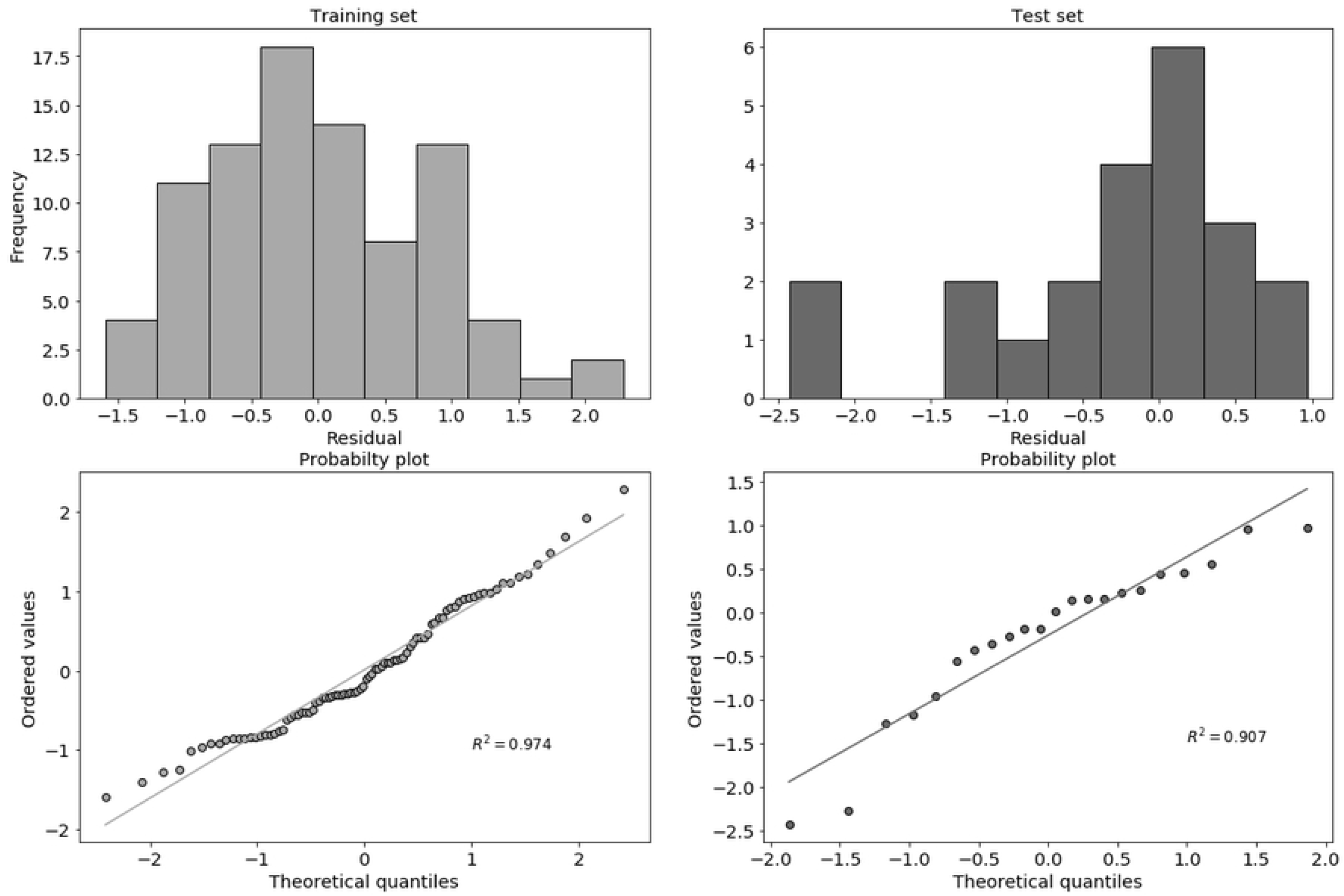
Histograms with the distibution of residuals, as predicted with Eq 5. The quantile plots, comparing to a normal distribution are also presented, for both the training and the test set. The regression line shows a near behavior to a normal distribution.

Observed and predicted activity values for both training and test sets are shown in Fig (4). The pIC_50_ values for the training set were calculated in a 10-fold cross validation step, thus the coefficient of determination in Fig (4) corresponds to *Q*^2^ — *LSO*. The test set, not used for the model construction, has a clear behavior near to the linear fit. The *R*^2^ for this external set is 0.71, above the typical threshold of 0.6. However, although a high value of both *Q*^2^ and *R*^2^ is required, it is not sufficient for the predictability estimation since these parameters just measure the linear correspondence between predicted and experimental values but not their 1:1 identity relationship [20]. Since there is not consensus in the establishment of an universal predictability criteria for QSAR modeling, one of the proposed practices is to calculate a set of parameters that could characterize the deviation from an ideal prediction, as suggested by Chirico *et al* [30] and Gramatica *et al* [31].

**Fig 4.**
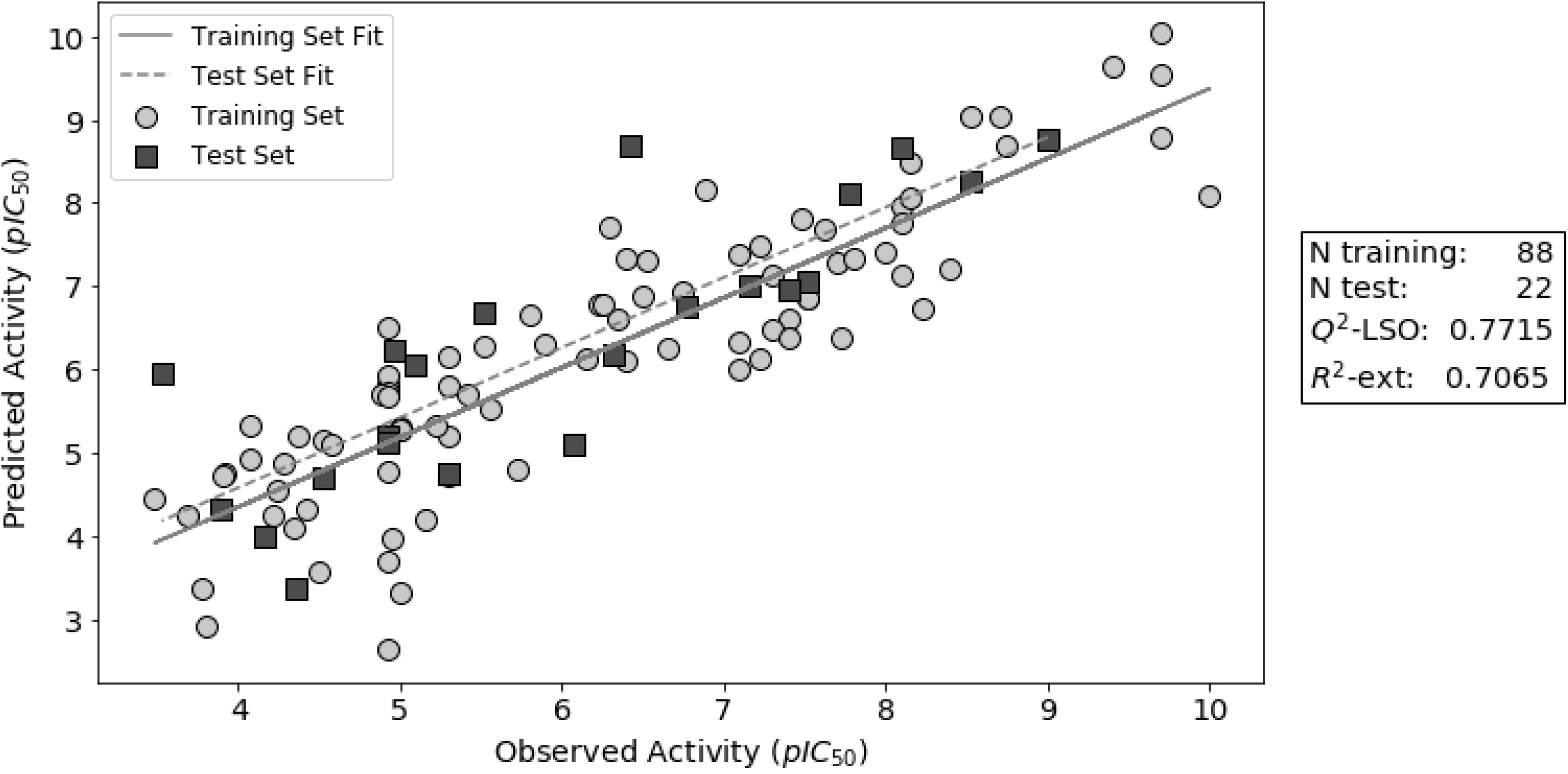
Regression plot for the results of predicted pIC_50_ values. The training set values shown were obtained in a 10-fold cross validation step. The coefficients of determination for both sets (*Q*^2^ — *LSO* and *R*^2^ — *ext*) are also presented. Continuous and dashed gray lines are the linear fits for training and test set, respectively.

The G&T criteria measure the agreement between experimental and predicted values [20,30,31]. These validation parameters were developed following the idea that the regression line for a predictive model should be the identity relation, *Ŷ_i_* — *Ŷ_i_*. Thus, the values of the G&T criteria measure the deviation of the least-squares line for the model from the identity straight line. Table 3 shows the results of these criteria, for the external evaluation used in this work, along with their acceptance thresholds as suggested by the authors. All the values are within the acceptable range, indicating a good agreement between the experimental information and the predictions of the model using the external test set.

**Table 3.**
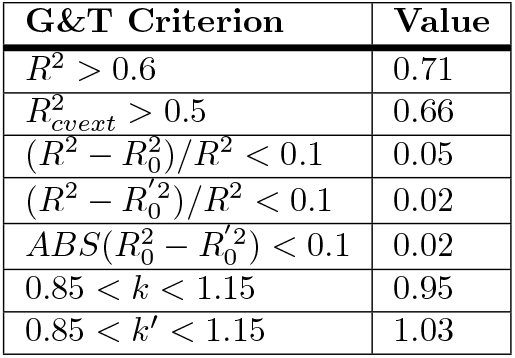
Golbraikh and Tropsha parameters and criteria for external validation calculated for the model

Applicability domain was defined using the leverage method, using both the training set and the test set. Williams plot for the dataset is presented in Fig (5). Because leverage is a projection of the distance from the training set, the distribution of the molecules in the Williams plot is a representation of the chemical space covered by the model. Standardized residuals are distributed around the expected value of 0, as was shown previously, for both the training and test sets. It is interesting to note that most of the test molecules follow a distribution similar to those in the training set, and their residuals are inside the expected errors predicted for the training set. It is also remarkable that two molecules in the training set and three in the test set display leverage values higher than the calculated limit. In these regions, any prediction made by the model is considered an extrapolation and its reliability is low.

**Fig 5.**
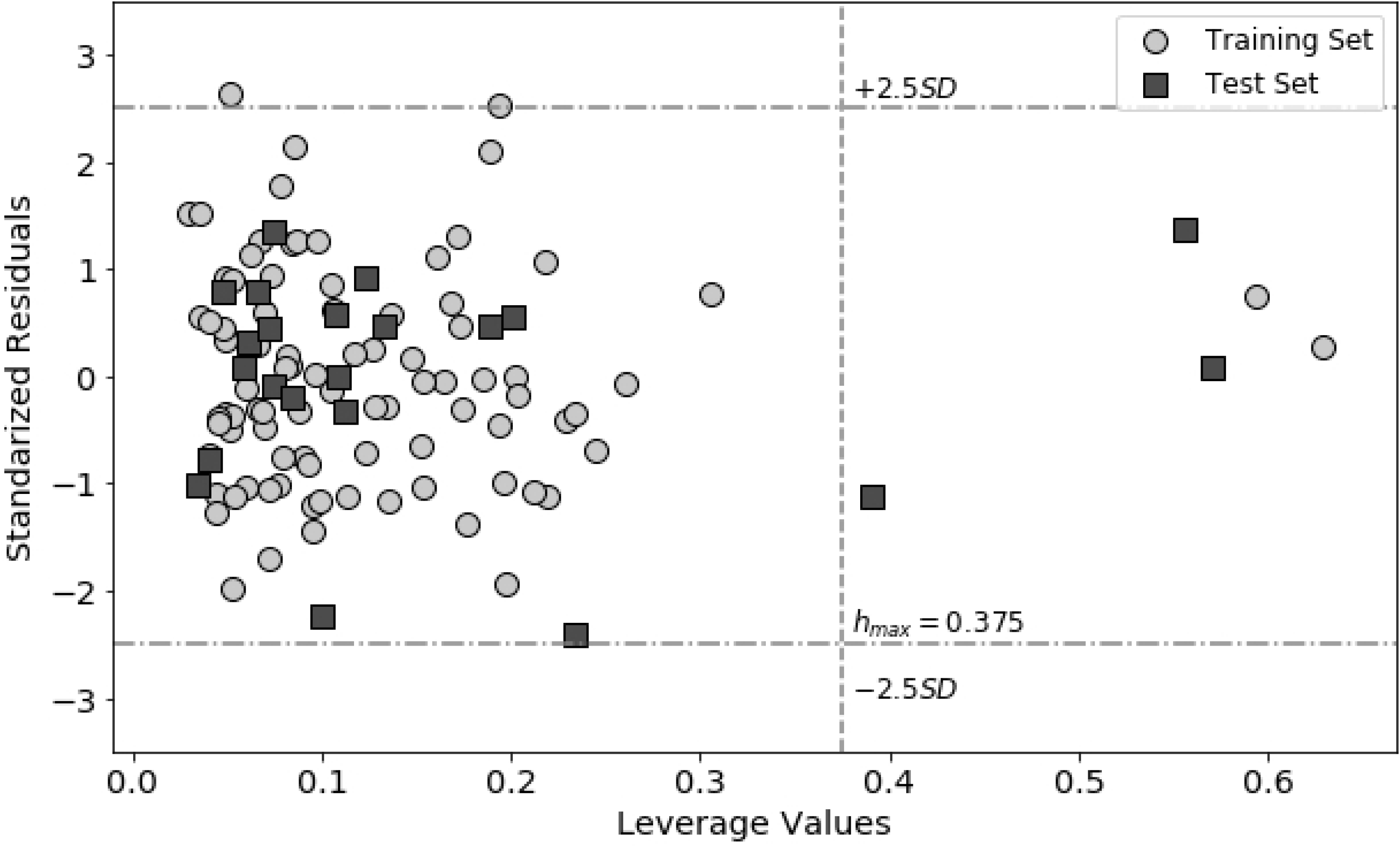
Williams plot for the applicability domain definition, using the leverage method.

Most of the molecular descriptors shown in Table 1 are related to potential intermolecular interactions. To rationalize the binding recognition process of cruzain inhibitors, based on the analysis of the molecular descriptors obtained in the QSAR model, molecular docking simulations were performed. PLIF histograms in Fig (6) summarize these results from the database of bound conformations. Fig (6A) shows interactions involving atom pairs between the protein and the ligand, whereas Fig (6B) summarizes surface contact interactions. These histograms show that hydrogen bond formation and polar contacts are predominant in the S1 subsite and near the catalytic site, whereas in S2 and S1’ subsites, hydrophobic contacts and π interactions are more favorable. Regarding with such interactions, molecular descriptor vsurf_JD8 is the hydrophobic integy moment (INTEraction enerGY) at −1.6 kcal/mol as defined by Cruciani *et al* [32]. Basically, the hydrophobic integy moment is the unbalance between the center of mass of the molecules and the hydrophobic regions. Thus, the descriptor may be related to the complementariety of inhibitors with the binding site, i.e. the ability to form hydrogen bonds or polar contacts with the catalytic site or the S1 subsite and hydrophobic or π interactions in the S2 or S1’ subsites.

**Fig 6.**
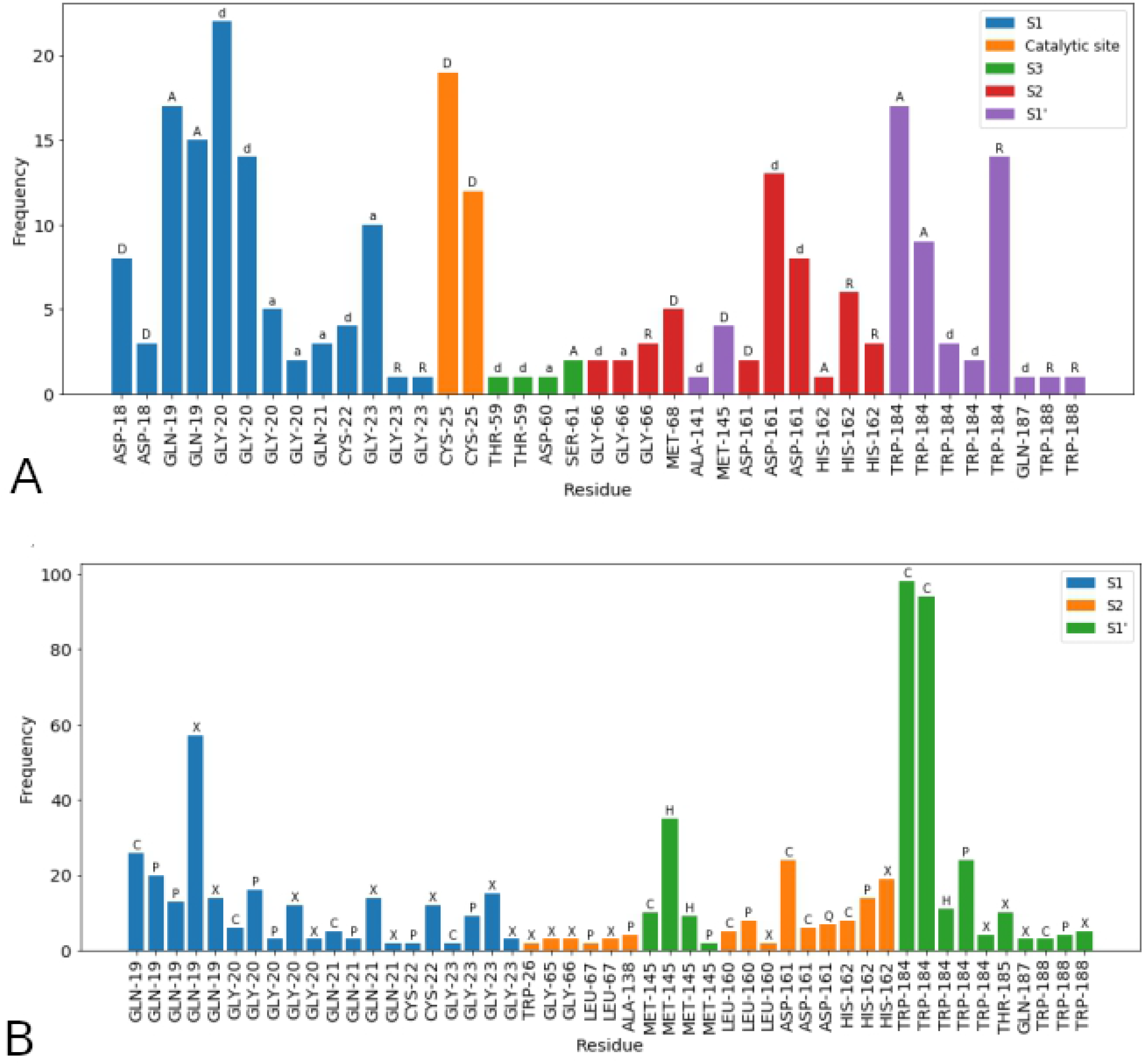
PLIF results for the docking calculation of the cruzain inhibitors in the dataset. A: PLIF histogram for potential contacts. The color of the bars represents the binding subsite in the cruzain. The code on the top of the bar is the kind of interaction: D, A, side chain hydrogen bond donors or acceptors; d, a, backbone hydrogen bond donors or acceptors, and R, arene or π interactions. B: PLIF histogram for surface contacts. The color of the bars represents the binding subsite in the cruzain. The code on the top of the bar is the kind of surface contact: H, hydrophobic; P, partial hydrophobic; Q, charged; X, other, and C total

Subdivided Van der Waals surface area descriptors are defined in terms of properties which can be divided into atomic contributions. In this case, partial charges and logP contributions take into consideration the total available surface area for certain types of electrostatic and hydrophobic contacts. Fig (7) shows the predicted conformations for some of the molecules in the set, along with their 2D representation depicting the partial charges and the atomic contributions to logP. PEOE_VSA_-1 and PEOE_VSA_-3 account the total surface area for atoms whose partial charges are in the ranges [—0.10, —0.05) and [—0.20, —0.15), respectively. Atoms with partial charges related to PEOE_VSA_-1 are often carbon atoms in aromatic rings and saturated chains. These molecular fragments bind to hydrophobic cavities, mainly in S2 and S1’ subsites and remarkably they have close contacts with TRP-184. On the other hand, partial charges accounted by PEOE_VSA_-3 are related to nitrogen-nitrogen containing groups, such as thiosemicarbazones, acylhydrazines and oxadiazoles. This partial charge is also associated with nitrile nitrogen, which is a chemical group present in the peptidomimetics, the most active compounds in the set. All these groups are frequently used as mimetics of the peptide bond since they can exert polar interactions required for the backbone recognition near the catalytic and S1 subsites.

**Fig 7.**
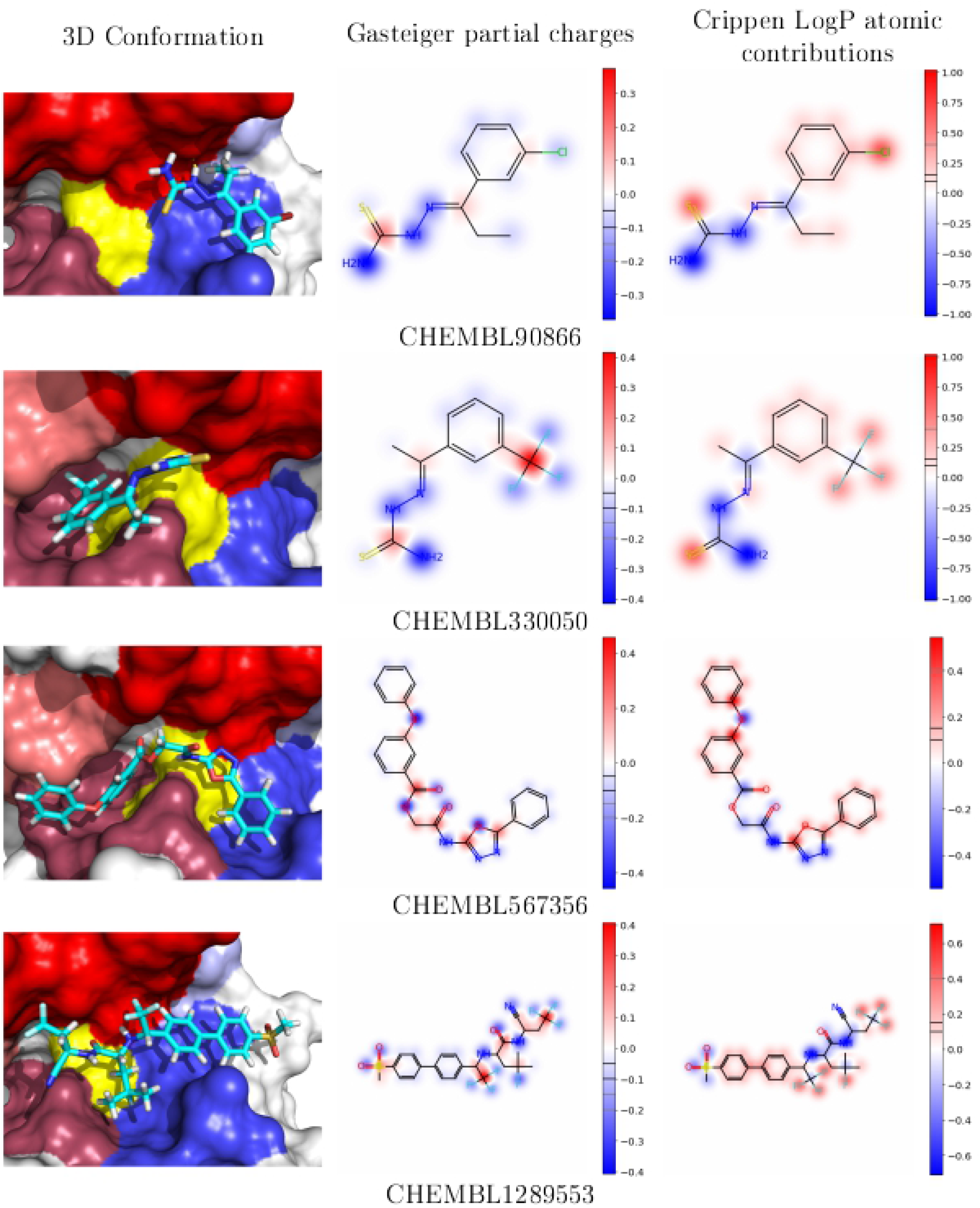
Binding conformations predicted by docking and visualization of descriptors related to partial charges and logP contributions. 2D depictions were generated as projections of their 3D conformations. The lines inside color bars are the ranges which contribute to the binned Van der Waals surface area descriptors in the QSAR model. In the 3D representation, cruzain subsites are shown in colors: yellow for the catalytic triad, red for S1 subsite, raspberry for the S2 subsite, deepsalmon for S3 subsite, tv_blue for S1’ subsite and lightblue for S2’ subsite (colors as defined by Pymol).

Regarding with surface descriptors, SLOGP_VSA4 and SLOGP_VSA9 measure the total surface area for logP atomic contributions in the ranges [0.1,0.15) and > 0.4, respectively. Most of the atomic fragments related to SLOGP_VSA4 are oxygen atoms in carbonyl groups directly attached to aromatic rings. However, the coefficient for this descriptor in the model equation is negative, indicating that this feature is unfavorable for biological activity. The last of the subdivided Van der Waals surface area descriptor takes into consideration mostly halogen atoms bound to aromatic or aliphatic groups. The most potent compounds in the data set are also rich in halogen-containing groups. Halogenated substituents are frequently used, among with other effects, to fulfill steric contacts into protein cavities, so they can exert a shape-complementary effect with the cruzain binding site, particularly in the well-defined S2 cavity and in the clefts formed by the S1 and S1’ subsites.

Volsurf descriptors are calculated from grids extended around the molecule, and then computing the interaction energy of this molecule with a probe on each of the grid points. DW12, EWmin1 and Wp5 are calculated using a water molecule as a probe, and thus are representative of polar interactions. For an energy isovalue of −3.0 kcal/mol, the field is representative of favorable polar and hydrogen bond donor-acceptor regions [32]. The total polar volume at this energy (Wp5) is positively correlated with biological activity, as can be deduced from its coefficient in the model equation. Futhermore, EWmin1 indicates that a lowest hydrophilic interaction energy is more favorable for cruzain inhibition. Fig (8) shows isosurfaces for the interaction fields at a level of −3.0 kcal/mol, with the same molecules as in Fig (7). It is clear from these representations that polar volumes extend around hydrogen bond donors and acceptors, mainly on those groups that mimic the peptide bond. Thus, these grid-based descriptors account for the ability of inhibitors to form hydrogen bonds in the binding site for the peptide bond recognition.

**Fig 8.**
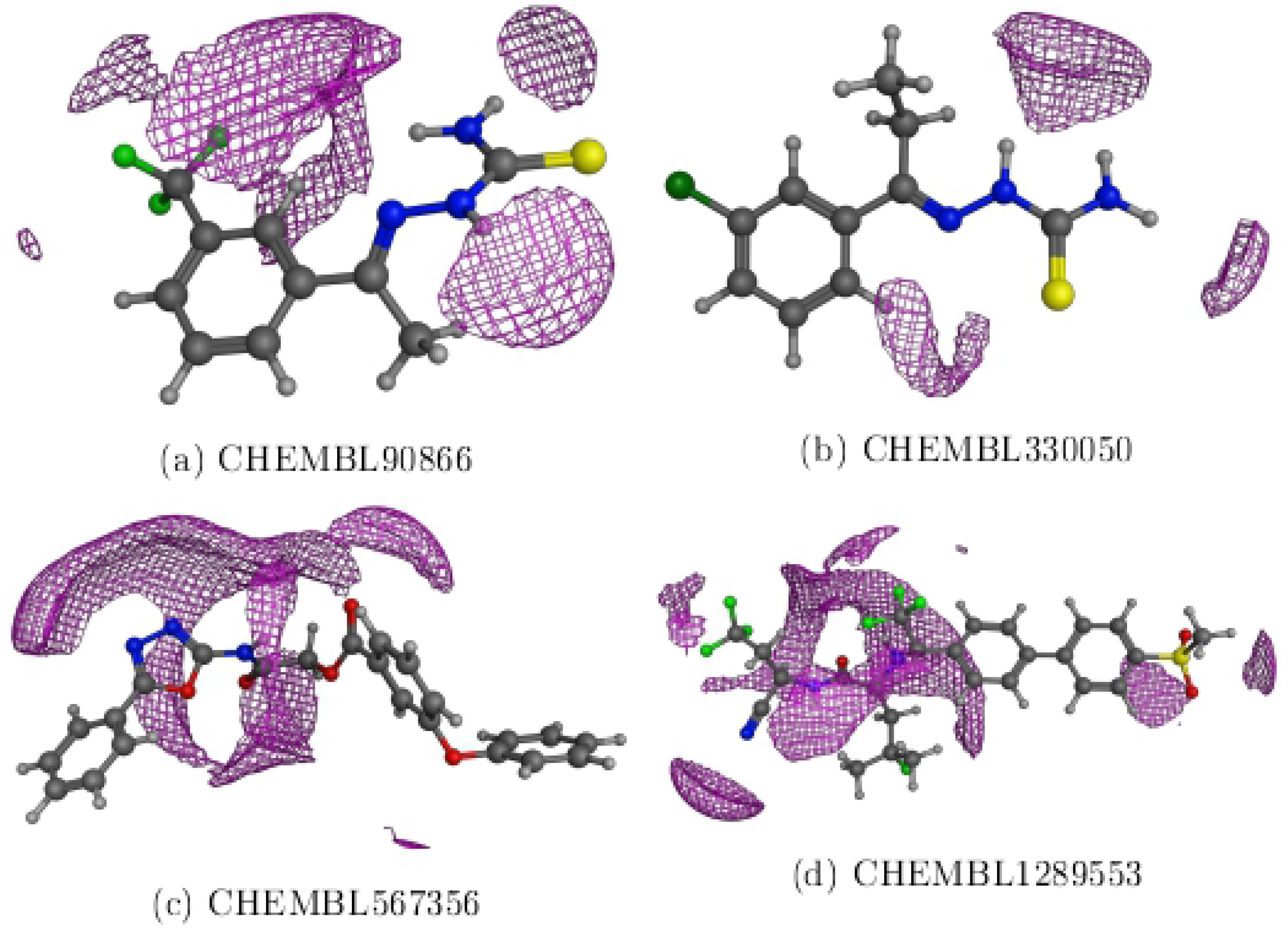
Polar surfaces at an isovalue of −3.0 kcal/mol. These interaction grids are calculated using a water molecule in every point and are indicative of polar interactions.

The interpretation provided above is based on the physical meaning of descriptors in terms of protein-ligand interactions. The model equation summarizes the presence of chemical fragments whose atoms meet the electrostatic and hydrophobic requirements for the binding into cruzain subsites but also their spatial distribution, as described by their integy moments and polar molecular fields. These requirements resemble a pharmacophore model that molecules within the applicability domain must meet to bind into the protein and exert its inhibitory effect.

In summary, we have presented a QSAR model with a well-defined endpoint, as described in methodology section for the criteria of data selection. The algorithm is unambiguously presented, which consists in the application of Eq 5 to calculate the predicted pIC_50_ for cruzain inhibition, given the required descriptors. The applicability domain is defined using the leverage method, and a limit value is also given for the reliability of predictions. The statistics for the goodness-of-fit, robustness and predictability were calculated and all of them fall within the generally accepted thresholds. Finally, a possible mechanistic interpretation of the model is proposed, in terms of intermolecular interactions. Thus, in this study, the five OECD principles for good practices in QSAR modeling are fulfilled. These principles are quality standards for QSAR developement, mainly in regulatory purposes. Under these criteria, our QSAR model is predictive and could be used in the search of new inhibitors or in the rational design of new compounds with this biological activity.

## Conclusion

A Quantitative Structure-Activity Relationship model was developed for the calculation of pIC_50_ values of cruzain inhibitors using multiple linear regression. The statistical parameters describing its performance agree with the general recommendations for QSAR modeling. In particular, the external validation demonstrates high predictability, since the calculated statistical parameters are above the recommended thresholds, considering its applicability domain. The molecular descriptors selected in the model equation are related to the potential formation of intermolecular interactions as shown in the binding modes calculated by docking. The linear equation integrates partial charge, hydrophobic potentials, and energy with spatial distribution and volume availability for polar interactions, indicating that there is a pharmacophoric-like recognition in the core of this QSAR model. The use and interpretation of this model could guide in the search, development and rational design of cruzain inhibitors as possible pharmacological treatment of Chagas disease.

## Supporting information

**S1 Table. Data set of cruzain inhibitors.** Table with the cruzain inhibitors, incluiding SMILES representation, activity values (pIC_50_), and calculated descriptors.

## Acknowledgments

K.M.-M. thanks DGAPA-UNAM (PAPIIT IN210518) and Instituto de Química, UNAM for finantial support. J.G.R.-J. thanks Biosen Institute for scholarship. Authors thank AutoDock Vina, Weka, Knime, RDKit, and Sci-Kit Learn developers for making machine learning and chemoinformatics tools freely available for academic purposes.

